# Basal ganglia contributions during the learning of a visuomotor rotation: effect of dopamine, deep brain stimulation and reinforcement

**DOI:** 10.1101/599308

**Authors:** Puneet Singh, Abhishek Lenka, Albert Stezin, Ketan Jhunjhunwala, Pramod Kumar Pal, Ashitava Ghosal, Aditya Murthy

## Abstract

It is commonly thought that visuomotor adaptation is mediated by the cerebellum while reinforcement learning is mediated by the basal ganglia. In contrast to this strict dichotomy, we demonstrate a role for the basal ganglia in visuomotor adaptation (error-based motor learning) in patients with Parkinson’s disease (PD) by comparing the degree of motor learning in the presence and absence of dopamine medication. We further show similar modulation of learning rates in the presence and absence of subthalamic deep brain stimulation. We also report that reinforcement is an essential component of visuomotor adaptation by demonstrating the lack of motor learning in patients with PD during the ON-dopamine state relative to the OFF-dopamine state in the absence of a reinforcement signal. Taken together, these results suggest that the basal ganglia modulate the gain of visuomotor adaptation based on the reinforcement received at the end of the trial.

## Introduction

Traditionally, visuomotor adaptation (error based) and reinforcement learning (reward based) are thought to occur through anatomically and functionally separate mechanisms (Doya, 1999, 2000; Hikosaka *et al.*, 2002; Haith & Krakauer, 2013; Taylor *et al.*, 2014). The mechanism underlying visuomotor adaptation is believed to minimize the differences between predicted and actual sensory feedback (Miall & Wolpert, 1996; Shadmehr & Krakauer, 2008; Shadmehr *et al.*, 2010). Reinforcement learning is considered to occur by selecting the motor commands that maximize reward or minimize punishment (Dam *et al.*, 2013; Wu *et al.*, 2014; Therrien *et al.*, 2016). Evidence for the independence of visuomotor adaptation and reinforcement learning is derived from experimental studies of visuomotor rotations that result in a recalibration of an internal forward model for error based learning during adaptation but not for reinforcement based learning (Izawa & Shadmehr, 2011). Visuomotor adaptation is also believed to be mediated by the cerebellum and reinforcement based motor learning by the basal ganglia (Doya, 2000). In support of this notion, patients with hereditary or acquired cerebellar damage have selective impairment in supervised (error-based) learning (Martin *et al.*, 1996; Maschke, 2003; Smith, 2005; Tseng *et al.*, 2007; Criscimagna-Hemminger *et al.*, 2010; Donchin *et al.*, 2012). However, they show no impairment during reinforcement learning of the same task (Izawa *et al.*, 2012; Therrien *et al.*, 2016).

In contrast, previous works on patients with Parkinson’s disease to identify the role of basal ganglia in modulating visuomotor adaptation have reported mixed results. While some studies showed no deficits in visuomotor adaptation (Marinelli *et al.*, 2009; Bédard & Sanes, 2011; Leow *et al.*, 2012; Semrau *et al.*, 2014), a different study has shown that dopaminergic medication modulates visuomotor adaptation (Mongeon *et al.*, 2013). Nevertheless, two potential shortcomings of these prior studies is that the effects reported were group wise effects and may have not been sensitive enough to reveal modulatory influences on visumotor learning. Further, the presence of dopaminergic effects per se, if any, does not directly implicate the basal ganglia in visuomotor adaptation since the cerebellum, as well as other diverse areas of the motor cortex that participate in motor learning, also receive independent dopaminergic projections from the ventral tegmental area (Ikai *et al.*, 1992; Melchitzky & Lewis, 2000).

Contrary to the conventional view that visuomotor adaptation is independent of reinforcement learning, the existence of bi-directional anatomical pathways between the cerebellum and the basal ganglia (Hoshi *et al.*, 2005; Bostan *et al.*, 2010) suggests that these structures are not necessarily separate information processing units. In particular, it has been demonstrated (Bostan *et al.*, 2010) that the cerebellum has a strong di-synaptic projection to the striatum through the thalamus, whereas the subthalamic nucleus has projections to the cerebellar cortex through the pontine nuclei (Bostan *et al.*, 2010). Consistent with these anatomical connections, some studies suggest that adaptation to visuomotor rotations involves a combination of different processes including reinforcement learning (Huang *et al.*, 2011; Nikooyan & Ahmed, 2015), as well as the use of explicit aiming strategies (Taylor *et al.*, 2014) but see (Herzfeld *et al.*, 2014; Leow *et al.*, 2016) for different views. In this context, a recent study showed how reinforcement and punishment differentially modulated the gain of learning in a visuomotor rotation task (Galea *et al.*, 2015), raising the hypothesis that the basal ganglia may modulate the sensitivity of cerebellum to errors thereby priming the cerebellum to weight its predictions or to update its internal model based on reinforcement received at the end of the trial.

To better understand the role of basal ganglia and understand the mechanisms, that is, whether basal ganglia makes an independent independent contribution to error based learning, or whether it modulates visuomotor adaptation through gain changes of error sensitivity, we manipulated the extent of dopamine (levodopa), assessed the effect of stimulation of subthalamic nucleus (STN) and reinforcement, visuomotor adaptation in patients with PD during several states: (i) with and without medication, (ii) with and without stimulation of the STN in patients who had undergone deep brain stimulation (DBS), (iii) in the presence and absence of a reinforcement signal.

## Materials and Methods

### Subjects

A total of 116 (64 patients and 52 healthy) individuals participated in this study. Patients were recruited from the neurology outpatient clinics and movement disorders services of the National Institute of Mental Health & Neurosciences, Bangalore, India. For Experiment 1, we recruited 20 patients with autosomal dominant cerebellar ataxia and 20 age-matched healthy controls. Assessment of the severity of ataxia was done by the International Cooperative Ataxia Rating Scale (ICARS). Further details about the ataxia patients’ characteristics and scores are shown in Table 1. For Experiment 2, we recruited 20 patients with idiopathic Parkinson’s disease (PD), and 20 age and gender matched healthy controls characteristics and scores are shown in Table 2. For Experiment 3, we recruited 12 idiopathic Parkinson’s disease patients with bilateral STN deep brain stimulation (DBS). Further details about the DBS patients’ characteristics and parameters of stimulation are shown in Table 3. For Experiment 4, we recruited 12 patients with idiopathic Parkinson’s disease (PD), and 12 age and gender matched healthy controls and details about the Parkinson’s disease patients are listed in Table 4. The diagnosis of PD was made as per the UK brain bank criteria (Hughes *et al.*, 1992). The forty healthy control subjects used in experiment 1 and experiment 2, were part of the data utilized in a previous paper for a separate purpose (Singh *et al.*, 2016). The section-III assessed motor symptoms of PD of the Unified Parkinson’s Disease Rating Scale (UPDRS-III) both during OFF-dopamine and ON-dopamine states and OFF medication with DBS ON and OFF. Mini-mental state examination (MMSE) was used to screen participants for cognitive impairment and patients with MMSE < 26 set as an exclusion criterion. All participants had normal or corrected to normal vision and no cognitive deficits. The handedness of subjects was tested by the modified Edinburgh Handedness Index (Salmaso & Longoni, 1985). The study was approved by the Indian Institute of Science ethics review board, and all the participants gave informed consent.

**Table 1:**
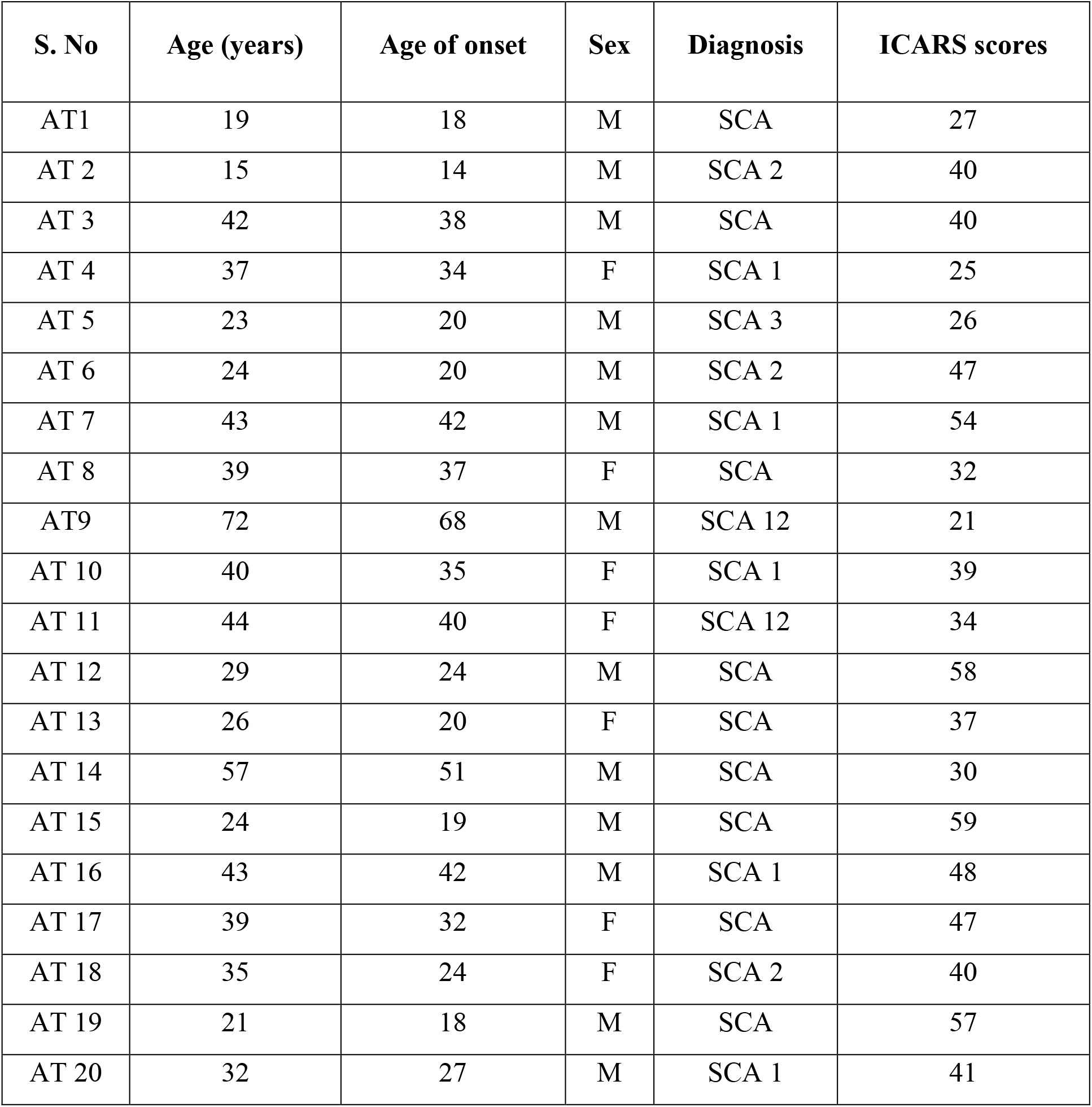
Demographics of cerebellar ataxia patients. AT = ataxia patients group; SCA = spinocerebellar ataxia types; ICARS = International Cooperative Ataxia Rating Scale.

**Table 2:** Demographics of Parkinson’s disease patients. PD = Parkinson disease patient group; H & Y Index = Hoehn and Yahr scale; UPDRS = Unified Parkinson’s Disease Rating Scale. The OFF-dopamine state was induced by withholding dopaminergic medication for at least 12 hours before the test. The ON-dopamine state was the best possible improvement after taking a supramaximal dose of Levodopa (usually 60-90 minutes after taking levodopa)

| S. No | Age | Age of onset | Sex | H & Y Index | UPDRS scores |  |
| --- | --- | --- | --- | --- | --- | --- |
|  |  |  |  |  | OFF-dopamine | ON-dopamine |
| PD1 | 64 | 53 | M | 1.5 | 12 | 03 |
| PD2 | 22 | 21 | M | 1.5 | 20 | 07 |
| PD3 | 40 | 37 | M | 02 | 35 | 04 |
| PD4 | 58 | 55 | M | 02 | 30 | 16 |
| PD5 | 60 | 52 | F | 02 | 25 | 11 |
| PD6 | 55 | 54.5 | F | 01 | 14 | 04 |
| PD7 | 68 | 60 | M | 02 | 19 | 13 |
| PD8 | 45 | 40 | F | 02 | 35 | 15 |
| PD9 | 40 | 36 | F | 01 | 30 | 11 |
| PD10 | 38 | 35.5 | M | 1.5 | 11 | 04 |
| PD11 | 56 | 50 | M | 2.5 | 44 | 31 |
| PD12 | 48 | 40 | F | 02 | 15 | 05 |
| PD13 | 58 | 49 | M | 02 | 19 | 11 |
| PD14 | 66 | 60 | F | 03 | 69 | 25 |
| PD15 | 49 | 33 | F | 2.5 | 30 | 18 |
| PD16 | 38 | 34 | M | 1.5 | 13 | 03 |
| PD17 | 66 | 56 | M | 1.5 | 22 | 05 |
| PD18 | 54 | 42 | M | 03 | 48 | 07 |
| PD19 | 70 | 61.5 | M | 1.5 | 24 | 14 |
| PD20 | 48 | 45 | M | 1.5 | 41 | 10 |

**Table 3:**
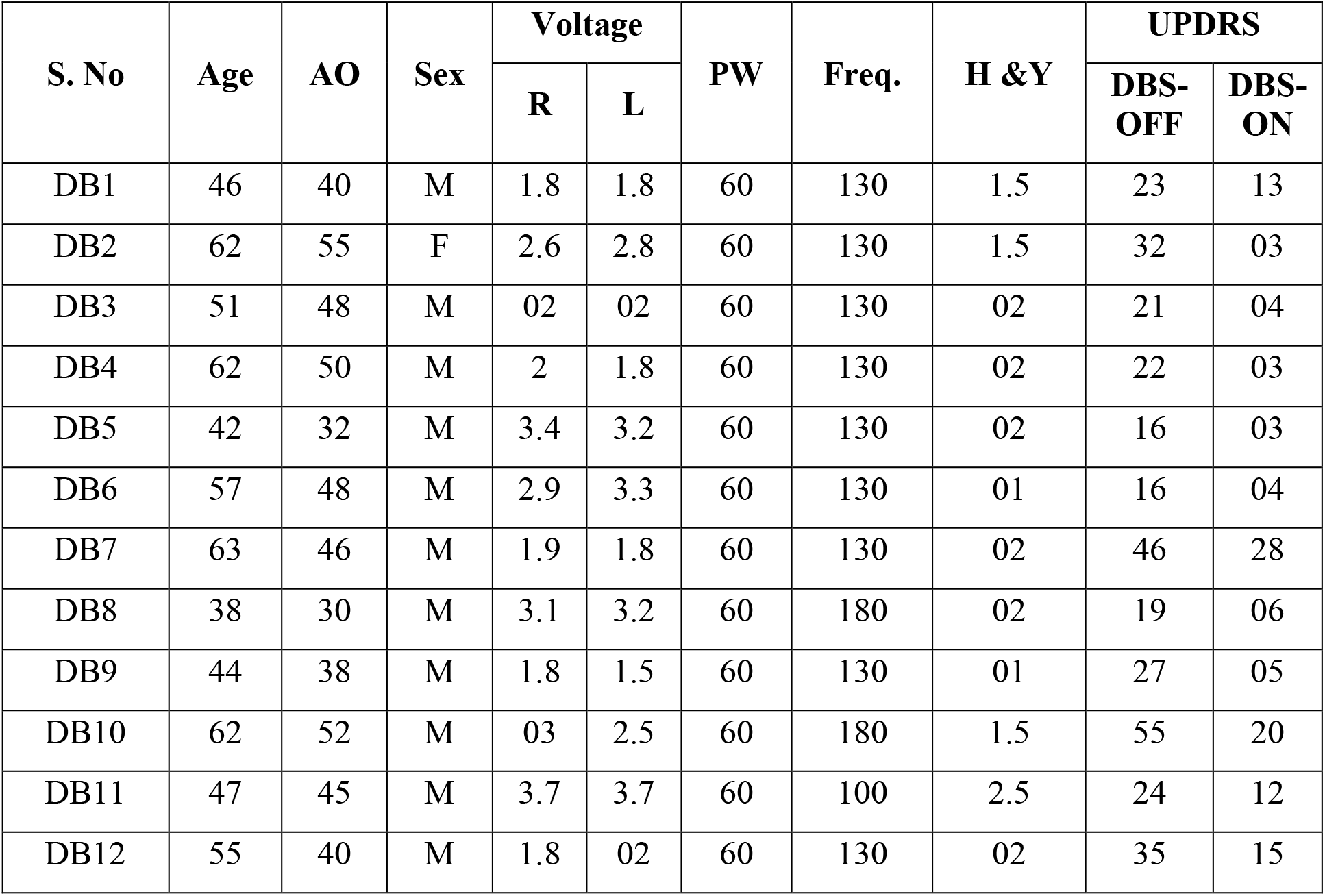
Demographics of Parkinson’s disease patient’s with deep brain stimulation. DB = deep brain stimulation patient group; AO = Age of onset; H & Y Index = Hoehn and Yahr scale; UPDRS = Unified Parkinson’s Disease Rating Scale. To test the impact of STN stimulation only, all patients remained OFF medication for a minimum period of 12 hours before the test.

**Table 4:** Demographics of Parkinson’s disease patients without reinforcement. NR = Parkinson patients without reinforcement signal group; H & Y Index = Hoehn and Yahr scale; UPDRS = Unified Parkinson’s Disease Rating Scale. The OFF-dopamine state was induced by withholding dopaminergic medication for at least 12 hours before the test. The ON-dopamine state was the best possible improvement after taking a supramaximal dose of Levodopa (usually 60-90 minutes after taking levodopa)

| S. No | Age | Age of onset | Sex | H &Y Index | UPDRS scores |  |
| --- | --- | --- | --- | --- | --- | --- |
|  |  |  |  |  | OFF-dopamine | ON-dopamine |
| NR1 | 60 | 56 | M | 1.5 | 23 | 13 |
| NR2 | 53 | 48 | M | 1.5 | 32 | 03 |
| NR3 | 56 | 49 | M | 02 | 21 | 04 |
| NR4 | 52 | 44 | M | 02 | 22 | 03 |
| NR5 | 57 | 54 | M | 02 | 16 | 03 |
| NR6 | 57 | 54 | M | 01 | 16 | 04 |
| NR7 | 56 | 50 | M | 02 | 46 | 28 |
| NR8 | 47 | 42 | M | 02 | 19 | 06 |
| NR9 | 56 | 52 | M | 01 | 27 | 05 |
| NR10 | 77 | 60 | M | 1.5 | 55 | 20 |
| NR11 | 65 | 57 | F | 2.5 | 24 | 12 |
| NR12 | 43 | 37 | F | 02 | 35 | 15 |

### Experimental setup

Participants sat on a chair with their hand placed on the front table as shown in Figure 1A. They looked straight ahead onto a monitor (refresh rate 60 Hz) that displayed both targets and the instantaneous hand cursor position while they moved the cursor in a horizontal plane. The experiment was performed using the Psychophysics Toolbox (in MATLAB) that displayed visual stimuli, sampled and stored the data and other behavioral parameters. Hand positions and joint angles were recorded (spatial resolution of 7.62 mm) using an electromagnetic position and orientation tracking device (Polhemus, LIBERTY, USA).

**Figure 1:**
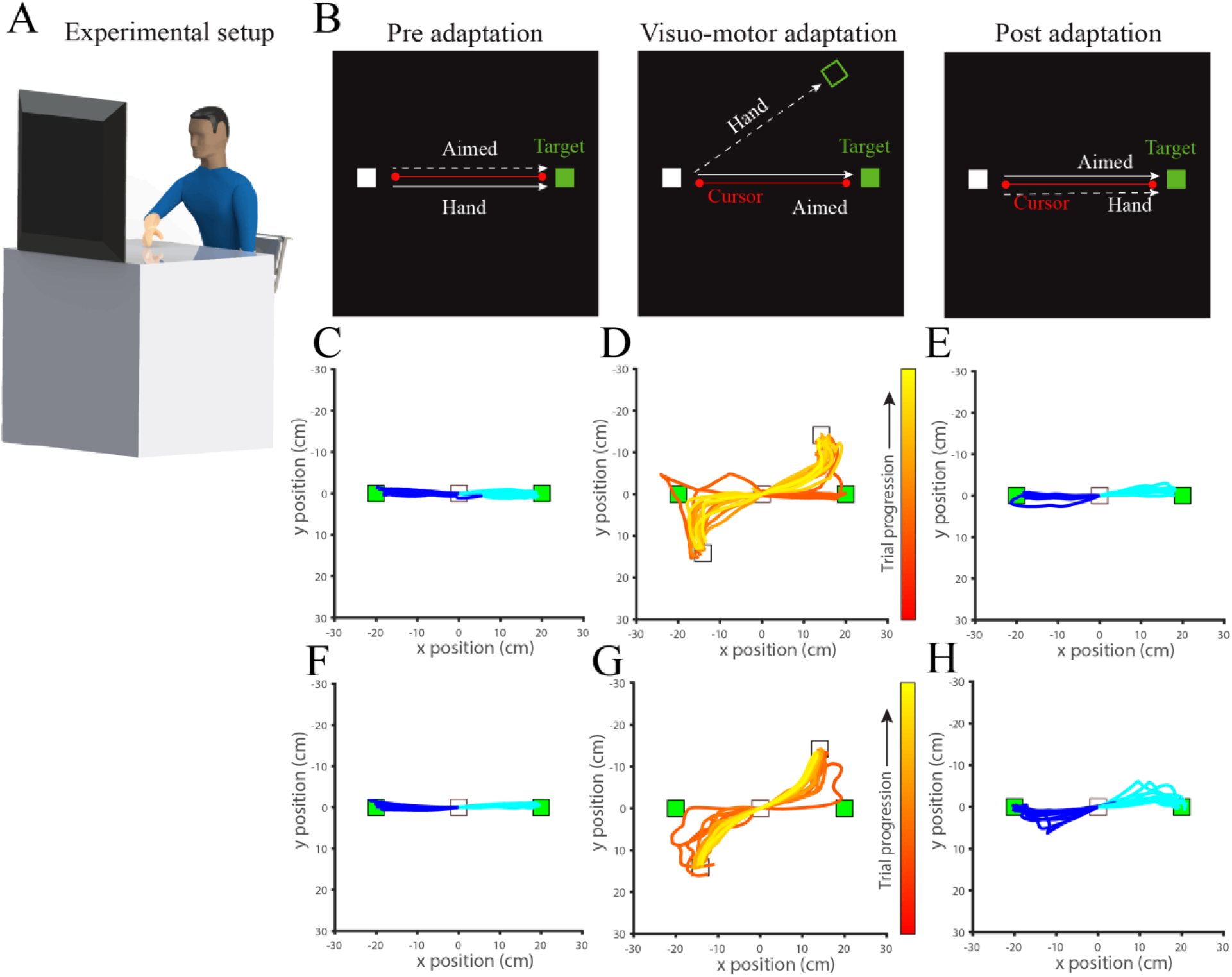
Experiment setup and design: (A) Subjects made point-to-point reaching movements to visual targets in 2 directions 20 cm away from the central start point in each trial. (B) Experiments were divided into a pre-adaptation (baseline), adaptation (visuomotor rotation) and post-adaptation (after-effect) epochs. (C) First five pre-adaptation trials from a patient subject showing baseline motor variability. (D) First ten visuomotor adaptation trials from the same patient subject showing the disturbed hand trajectory. (E) First five postadaptation trials from the same patient subject indicating the effect of adaptation. (F) First five pre-adaptation trials from a control subject showing baseline motor variability. (G) First ten visuomotor adaptation trials from the same control subject showing the disturbed hand trajectory. (H) First five post-adaptation trials from the same control subject indicating the effect of adaptation.

### Experimental paradigm

For all experiments, trials were divided into three phases – baseline or pre-adaptation, adaptation, and post-adaptation. Each trial started with the presentation of a square hand-fixation box (1.5 cm * 1.5 cm) at the center of the screen where the subject had to fixate the hand cursor. After successful hand-fixation, a square target (1.5 cm * 1.5 cm) was displayed randomly in any one of 2 locations 20 cm away from the central hand-fixation box. The subject moved their hand to the target only after the hand-fixation box disappeared. All subjects performed ~10 practice trials before the experimental session. Subjects performed the experimental paradigm with 100 trials per session, with a typical session lasting for 15-20 minutes. The instruction was to reach the target as fast as they could, and the maximum time window to complete the 20 cm movement was 2000 ms. The participants were not informed about the perturbation. No constraint of movement velocity was enforced or instructed since there was variability across the subjects, across disease state (Ataxia, PD, PD with DBS) and even between subject medication state (ON or OFF medication). During the visuomotor perturbation, the cursor movement was rotated according to equation (1)

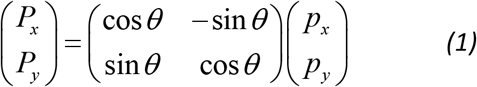

where *P_x_*, *P_y_* correspond to the position of the cursor, *p_x_*, *p_y_* correspond to the actual position of the hand and *θ* (−45°) denotes the perturbation angle about the center of work space. This perturbation led to a trajectory error that was gradually compensated over the course of many trials till the hand trajectory straightened again. In Experiments 1, 2 and 3, when subjects reached the target, they received a reinforcement signal consisting of an auditory tone presented at 900 Hz for 300 ms. The latency between acquiring the target and the reinforcement signal was ~20 ms. In Experiment 4, subjects did not receive any reinforcement signal. This was done to test the role of reinforcement on learning. The target disappeared after the completion of the trial, and participants had to return to the home position without any time constraints while the perturbation was still on. To ensure participants returned to the same starting position, the home position was always fixed and marked on the table.

### Quantifying learning

The error was calculated as the maximum perpendicular distance of the hand trajectory from the straight line joining the central to hand-fixation box to the target location. The error, denoted by f (n), is related to the trial number by the following equation.

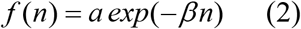

The above equation represents a first-order learning process and *β* represents the natural learning rate for a subject. To compute the population learning in perturbation trials, errors were fitted with an exponential fit using a robust least squares method.

### Statistical analysis

The data were assessed for normality using Lilliefors test. For pairwise comparisons between the groups, a two-tailed t-test was performed if the data was normally distributed otherwise Wilcoxon signed-rank test was used. Comparison of two independent groups was made using a two-sample t-test was used for normally distributed data; otherwise a Wilcoxon rank-sum test was used. Furthermore, to compute the effect size we used the Cohen’s d test as well as report 95% confidence intervals when parametric tests were used. All the correlational analyses were performed using Pearson’s correlation method.

## Results

### Visuomotor adaptation in patients with degenerative cerebellar disease (Experiment 1)

To confirm the role of the cerebellum in visuomotor adaptation, we assessed motor learning in patients with autosomal dominant cerebellar ataxia (n=20) and compared their learning with age and gender matched healthy controls (n=20) (Figure 1A & 1B). Both groups performed point-to-point reaching movement in a visuomotor adaptation task where the stimulus randomly alternates in two opposite directions. A perturbation was introduced by rotating the cursor by 45 degrees from the hand trajectory. Overall, the pattern of trajectories at the baseline (without the visuomotor perturbation) showed nearly straight trajectory across both groups but showed strongly curved trajectories in the presence of a visuomotor perturbation (Figure 1C & 1D). Consistent with previous literature, the curved trajectories gradually become straighter with practice over the course of about sixty trials in healthy controls but not in patients with cerebellar ataxia (Figure 2A). The reduction in maximum error (equation (2)) was used as a metric to quantify the learning rate for each subject. We observed that the mean learning rate for the cerebellar ataxia group (mean learning rate = 0.0071 ± 0.002, 95% CI 0.002 - 0.012) was significantly lower than the mean learning rate for the control group (mean learning rate = 0.023 ± 0.003, 95% CI 0.016 - 0.03) (Figure 2B; p = 3.3e-4, t (38) = 3.94, Cohen’s d = 1.24). We also compared the mean errors in perturbation trials, as opposed to the learning rates, to confirm the same result. The mean errors for the cerebellar ataxia group (mean errors = 7.11 ± 0.92, 95% CI 5.18 - 9.04) was significantly higher than the mean errors for the control group (mean errors = 4.24 ± 0.45, 95% CI 3.30 - 5.19) (Figure 2C; p = 0.0081, t (38) = 2.79, Cohen’s d = 0.88). Consequent to the absence of any overt motor learning, the cerebellar ataxia group showed no after-effect (post-adaptation; mean errors = −2.29 ± 0.87, 95% CI −4.11 - −0.47). The mean error during the first five trials post-adaptation was used to quantify the aftereffect. However, the control group showed the characteristic, albeit weak after-effect in the opposite direction when the learned visuomotor perturbation was turned off (mean errors = −6.22 ± 0.53, 95% CI −7.32 - −5.12) (Figure 2D; p = 4.21e-4, t (38) = 3.86, Cohen’s d = 1.22), This after-effect error reverted to the baseline levels typically within twenty trials.

**Figure 2:**
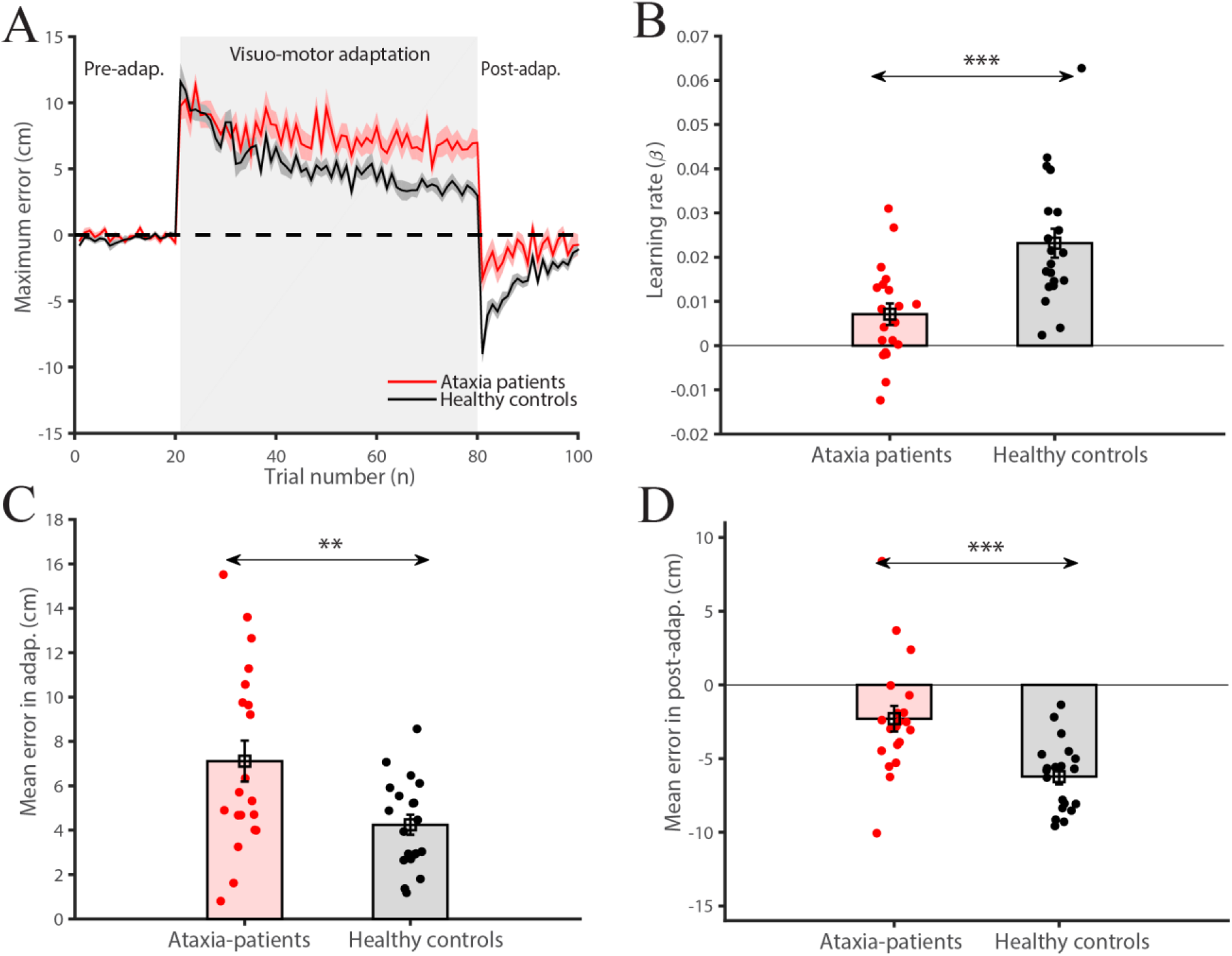
Ataxia patients in visuomotor adaptation. (A) The maximum error in pre-adaptation, visuomotor adaptation, and post-adaptation across Ataxia patients (n=20) and healthy controls. Red indicates Ataxia patients and black indicates healthy controls. (B) Learning rate differences in Ataxia patients (red) and healthy controls (black) (n=20) reveal faster learning in healthy controls. (C) Averaged mean errors in Ataxia patients (red) and healthy controls (black) (n=20) reveal higher error magnitude in Ataxia patients. (D) Averaged mean errors in Ataxia patients (red) and healthy controls (black) in post-adaptation revealed higher retention in healthy subjects. Solid lines indicate mean and error bars or shaded areas are SEM (*P < 0.05; **P < 0.005; ***P < 0.0005).

### Visuomotor adaptation in patients with Parkinson’s disease (PD; Experiment 2)

We used the performance of patients with PD and healthy controls in order to assess the contribution of basal ganglia during adaptation. We trained 20 patients with PD, and an equal number of age and gender matched healthy controls on the same two directions visuomotor rotation task as patients with cerebellar ataxia. To test the impact of dopaminergic medications, all patients with PD were tested in two sessions: (i) during the OFF-dopamine state, and (ii) during the ON-dopamine state. The OFF-dopamine state was induced by withholding dopaminergic medication for at least 12 hours before the test. The ON-dopamine state was the best possible improvement after taking a supramaximal dose of Levodopa (usually 60-90 minutes after taking levodopa). To avoid confounds due to the transfer of learning between sessions, the order of testing between ON and OFF medication was counterbalanced.

Overall, the pattern of trajectories at baseline was nearly straight across the groups and showed strongly curved trajectories in the presence of the visuomotor perturbation (Figure 3A). The curved trajectories gradually became straighter with practice over the course of about sixty trials in controls and patients with PD in the ON-dopamine state but not in the case of PD patients during the OFF-dopamine state. As before, the reduction in maximum error (equation (2)) was used as a metric to quantify the learning rate for each subject. The mean learning rate for PD patients in the OFF-dopamine state (mean learning rate = 0.005 ± 0.002, 95% CI 0.001 - 0.010) was significantly less than the mean learning rate for the same patients during the ON-dopamine state (mean learning rate = 0.020 ± 0.001, 95% CI 0.016-0.023) (Figure 3B; p = 1.8e-06, t (19) = 6.77, Cohen’s d = 1.51). There was no difference in the mean learning rate between the patients during ON-dopamine and healthy control group (mean learning rate = 0.022 ± 0.002, 95% CI 0.016 - 0.027) (Figure 3B; p = 0.56, t (38) = 0.582, Cohen’s d = 0.18). We also compared the mean errors in perturbation trials between OFF and ON medication groups (Figure 3C; OFF-dopamine mean errors = 6.91 ± 0.77, 95% CI 5.30 – 8.52; ON-dopamine mean errors = 4.64 ± 0.47, 95% CI 3.66 – 5.62; p = 0.0011, t (19) = 3.85, Cohen’s d = 0.86) and between ON-dopamine and healthy control group (control mean errors = 4.56 ± 0.37, 95% CI 3.78 – 5.33) (Figure 3C; p = 0.89, t (38) = 0.132, Cohen’s d = 0.04), which reconfirms the previous result, albeit using the mean errors as opposed to learning rates. As a consequence of minimal motor learning, patients OFF-dopamine showed much smaller after-effects (first five trials in post-adaptation) (Figure 3D; OFF-dopamine mean errors = −1.59 ± 0.62, 95% CI −2.88 – −0.31; ON-dopamine mean errors = −4.21 ± 0.55, 95% CI −5.38 – −3.05; p = 0.002, t (19) = 3.51, Cohen’s d = 0.79). In contrast, patients ON-dopamine demonstrated a classic after-effect comparable to controls when the learned visuomotor perturbation was turned off (Figure 3D; control mean errors = - 5.34 ± 0.62, 95% CI −6.64 – −4.04, p = 0.18, t (38) = 1.35, Cohen’s d = 0.42). To control for the order in which patients were trained on the task, we separated the patient group into those subjects that performed the task first from those subjects who performing the task second. This was done for the OFF-dopamine and the ON-dopamine subjects. As illustrated in (Figure 3E and 3F) the main results remain the same independent of order.

**Figure 3:**
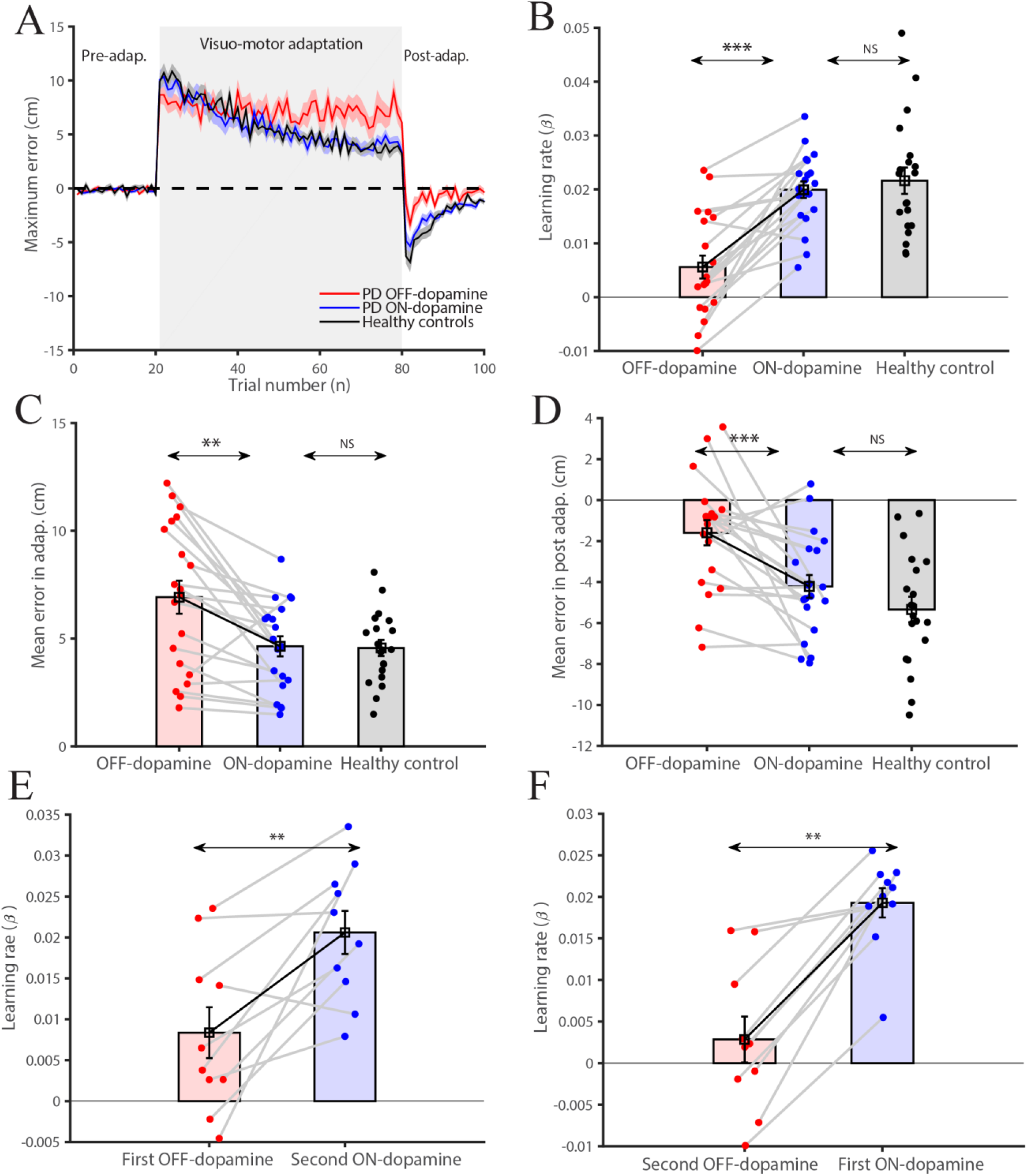
Parkinson’s patients in visuomotor adaptation (A) Maximum error in preadaptation, visuomotor adaptation, and post-adaptation in across Parkinson’s patients with medicine differences (n=20). Red indicates OFF-medicine; blue indicates ON-medicine and black indicate healthy controls. (B) Learning differences in the OFF-dopamine (red) and ON-dopamine (blue) (n=20), reveal faster learning in the ON-dopamine and show no differences in ON-dopamine and healthy controls. (C) Averaged mean errors in OFF-dopamine (red), ON-dopamine (blue) and healthy controls (black) (n=20) reveal higher error magnitude in OFF-dopamine condition. (D) Similarly, averaged mean errors in OFF-dopamine (red), ON-medicine (blue) and healthy controls (black) in post-adaptation revealed low retention in OFF-medicine condition. (E) Learning differences in the First OFF-dopamine (red) and Second ON-dopamine (blue) (n=10), reveal faster learning in the ON-dopamine. (F) Learning differences in the First ON-dopamine (blue) and Second OFF-dopamine (red) (n=10), reveal faster learning in the ON-dopamine.

### Visuomotor adaptation in patients with deep brain stimulation (STN-DBS; Experiment 3)

Although the previous result indicates a role of dopamine in visuomotor adaptation, the effects of oral levodopa formulation are not specific to basal ganglia but may involve projections directly to the cerebellum via the ventral tegmental area (VTA) (Ikai *et al.*, 1992, 1994). To demonstrate the role of basal ganglia in motor learning, a direct manipulation of structures in basal ganglia is necessary. To pursue this, we recruited patients with PD who had undergone deep brain stimulation (DBS) of the subthalamic nucleus (STN) to manipulate the basal ganglia (n=12). To test the impact of STN stimulation only, all patients remained OFF medication for a minimum period of 12 hours before the test. All patients were tested in two sessions: OFF-DBS and ON-DBS, with the order of the two sessions being counterbalanced across subjects.

Once again to quantify the error, we used the maximum error along the trajectory (Figure 4A). The reduction in maximum error (equation (2)) was used as a metric to quantify the learning rate for each subject. We observed that the mean learning rate for the OFF-DBS group (mean learning rate = −0.003 ± 0.002, 95% CI −0.008 – 0.003) was significantly less than the mean learning rate for the ON-DBS group (mean learning rate = 0.017 ± 0.003, 95% CI 0.009 – 0.024) (Figure 4B; p = 5.97e-06, t (11) = 8.07, Cohen’s d = 2.33). We did not observe any difference in the learning rates between ON-DBS and control groups (mean learning rate = 0.023 ± 0.004, 95% CI 0.013 – 0.034) (Figure 4B; p = 0.29, t (22) = 1.07, Cohen’s d = 0.43). Likewise, the mean errors in perturbation trials were significantly larger in OFF-DBS compared to ON-DBS groups (Figure 4C; OFF-DBS mean errors = 6.89 ± 0.60, 95% CI 5.56 – 8.22; ON-DBS mean errors = 4.90 ± 0.63, 95% CI 3.51 – 6.28; p = 0.06, t (11) = 2.07, Cohen’s d = 0.60), whereas ON-DBS and healthy control group mean errors were not significant different (Figure 4C; control mean errors = 4.09 ± 0.54, 95% CI 2.88 – 5.30; p = 0.34, t (22) = 0.96, Cohen’s d = 0.39). Interestingly, we observed differences in the after effects in DBS patients that was not observed in their learning abilities. In contrast to the adaptation period, the OFF-DBS and ON-DBS groups showed no significant difference in their after-effects, (Figure 4D; OFF-DBS mean errors = −1.03 ± 0.62, 95% CI −2.39 – 0.32; ON-DBS mean errors = −2.93 ± 0.66, 95% CI −4.40 – −1.46; p = 0.09, t (11) = 1.80, Cohen’s d = 0.52). In contrast, the healthy control group showed a robust after-effect relative to ON-DBS, that was not observable in their learning abilities (Figure 4D; control mean errors = – 6.31 ± 0.79, 95% CI −8.05 – −4.56; p = 0.003, t (22) = 3.26, Cohen’s d = 1.33). To control for the order in which patients were trained on the task, we separated the patient group into those subjects that performed the task first from those subjects who performing the task second. This was done for the OFF-DBS and the ON-DBS subjects. As illustrated in Figure 4E and 4F the main results remain the same independent of order.

**Figure 4:**
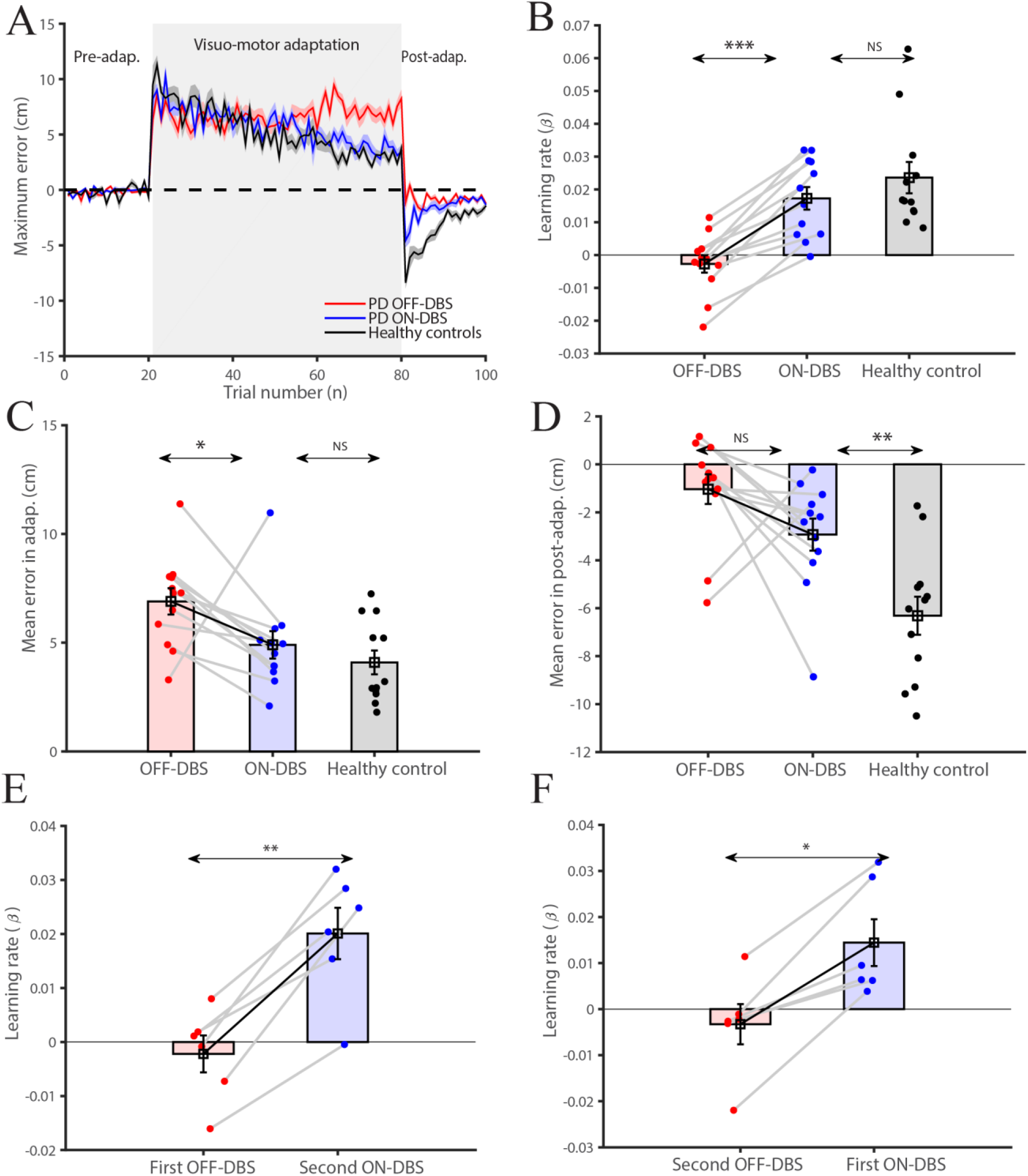
Parkinson’s patients with subthalamic deep brain stimulation (A) Maximum error during pre-adaptation (baseline), visuomotor adaptation, and post-adaptation as a function of the trial number in PD patients with and without DBS relative to healthy controls. Red indicates OFF-DBS, blue indicates ON-DBS and black indicates healthy controls. The learning curves are an average across the population (n=12). The shaded area indicates the corresponding SE shown. (B) Learning rates in the OFF-DBS (red) and ON-DBS (blue) conditions relative to healthy controls (black; n=12 for each group). (C) Averaged mean errors in OFF-DBS (red), ON-DBS (blue) and healthy controls (black) (n=20) reveal higher error magnitude in OFF-DBS condition. (D) Similarly, averaged mean errors in OFF-DBS (red), ON-dopamine (blue) in post-adaptation revealed low retention in OFF-DBS and ON-DBS conditions in comparison to healthy controls. (E) Learning differences in the First OFF-DBS (red) and Second ON-DBS (blue) (n=6), reveal faster learning in the ON-DBS. (F) Learning differences in the First ON-DBS (blue) and Second OFF-DBS (red) (n=6), reveal faster learning in the ON-dopamine.

### Assessing the role basal ganglia in visuomotor adaptation: the role of reinforcement (Experiment 4)

Although, the level of dopamine and STN stimulation appears to modulate the rate of adaptation in what is traditionally thought to be driven by error, we examined whether this modulation was a consequence of motivation provided by the auditory feedback, which was a secondary reinforcement signal given to subjects following successful completion of the trial (i.e., the cursor reaching the target location). It is important such reinforcement is all or none and given at the end of the trial and is different from the error that gradually reduces over time during the course of the trial. To test this hypothesis, we trained a new set of PD patients (n=12) during the OFF-dopamine and ON-dopamine states, and an equal number age and gender matched healthy controls with the same visuomotor rotation task but in the absence of auditory reinforcement feedback. Similar to the previous experiment, the pattern of trajectories in the baseline condition showed nearly straight trajectory across groups but showed strongly curved trajectories in the presence of a visuomotor perturbation (Figure 5A). Surprisingly, in the absence of reinforcement, the median learning rate for the patients in the OFF-dopamine state (median learning rate = 0.008 ± 0.003) was similar to the median learning rate for the same patients during the ON-dopamine state (Figure 5B; median learning rate = 0.018 ± 0.002, p = 0.064, signed rank =15). We also observed a difference in the median learning rate between the patients during ON-dopamine and healthy control group in absence of reinforcement (median learning rate = 0.025 ± 0.005, 95% CI 0.015 – 0.039) (Figure 5B; p = 0.04, rank sum= 115). Likewise, we observed no differences the mean errors in perturbation trials in OFF and ON-dopamine without reinforcement groups (Figure 5C; OFF-dopamine mean errors = 5.23 ± 0.62, 95% CI 3.86 – 6.61; ON-dopamine mean errors = 4.50 ± 0.49, 95% CI 3.42 – 5.58; p = 0.18, t (11) = 1.41, Cohen’s d = 0.41), whereas ON- dopamine and healthy control without reinforcement group mean errors were significant different (Figure 5C; control mean errors = 3.16 ± 0.40, 95% CI 2.27 – 4.05; p = 0.04, t (22) = 2.11, Cohen’s d = 0.86). Similarly, to the adaptation period the OFF-dopamine and ON-dopamine a without reinforcement groups showed no significant difference in their aftereffects, (Figure 5D; OFF-dopamine median errors = −2.39 ± 0.58, 95% CI −3.95 – −1.39; ON-dopamine median errors = −2.40 ± 0.79; p = 0.10, sign rank = 60). Similarly, the healthy control group also did not show an after-effect without reinforcement (Figure 5D; control median errors = −3.34 ± 0.72; p = 0.70, rank sum = 157).

**Figure 5:**
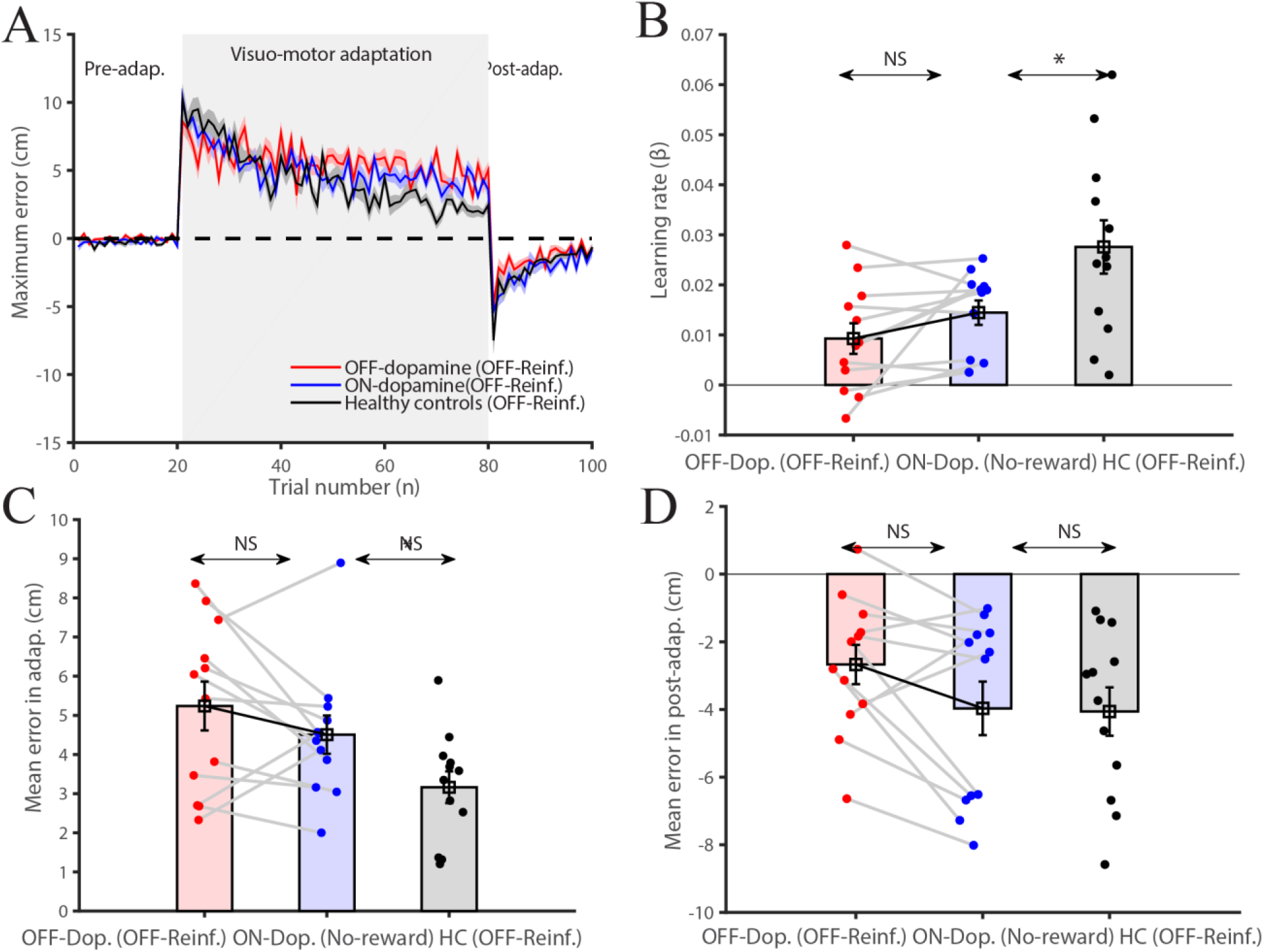
Parkinson’s patients without reinforcement in visuomotor adaptation (A) Maximum error in pre-adaptation, visuomotor adaptation, and post-adaptation across Parkinson’s patients with medicine differences plus off reinforcement (n=12). Red indicates OFF-dopamine with off reinforcement, blue indicates ON-dopamine with off reinforcement and black indicates healthy controls with off reinforcement. (B) Learning differences in the OFF-dopamine with off reinforcement, (red) and ON-medicine with off reinforcement (blue) reveal no differences in ON and OFF-dopamine and also reveal faster learning in the healthy controls. (C) Averaged mean errors in OFF-dopamine with off reinforcement (red), ON-dopamine with off reinforcement (blue) and healthy controls with off reinforcement (black) (n=20) reveal no differences error magnitude in OFF and ON-dopamine with off reinforcement condition. (D) Similarly, averaged mean errors in OFF-dopamine (red), ON-medicine (blue) in post-adaptation revealed less retention in OFF-dopamine, ON-dopamine, and healthy controls.

Furthermore, when we compared healthy control subjects with and without reinforcement (Figure 6A), the mean learning rate in the presence of reinforcement (mean learning rate = 0.023 ± 0.004, 95% CI 0.013 – 0.034) was similar to the mean learning rate without reinforcement (Figure 6B; mean learning rate = 0.027 ± 0.005, 95% CI 0.016 – 0.039, p = 0.58, t (22) = 0.55, Cohen’s d = 0.22). Likewise, we observed no differences the median errors in perturbation trials in OFF and ON-dopamine without reinforcement (Figure 6C; with reinforcement median errors = 3.07 ± 0.54, 95% CI 2.88 – 5.30; without reinforcement median errors = 3.46 ± 0.40, 95% CI 2.27 – 4.50; p = 0.40, rank sum = 165). Interestingly, we observed differences in the after effects in healthy control with and without reinforcement. (Figure 6D; with reinforcement mean errors = −6.31 ± 0.79, 95% CI −8.06 – −4.56; without reinforcement mean errors = −4.06 ± 0.71, 95% CI −5.63 – −2.48; p = 0.046, t (22) = 2.11, Cohen’s d = 0.86). In summary, we observed that the absence of reinforcement interestingly abolished the difference between rate of adaptation we had previously observed between patients ON versus OFF medication as well as those ON versus OFF DBS of the STN.

**Figure 6:**
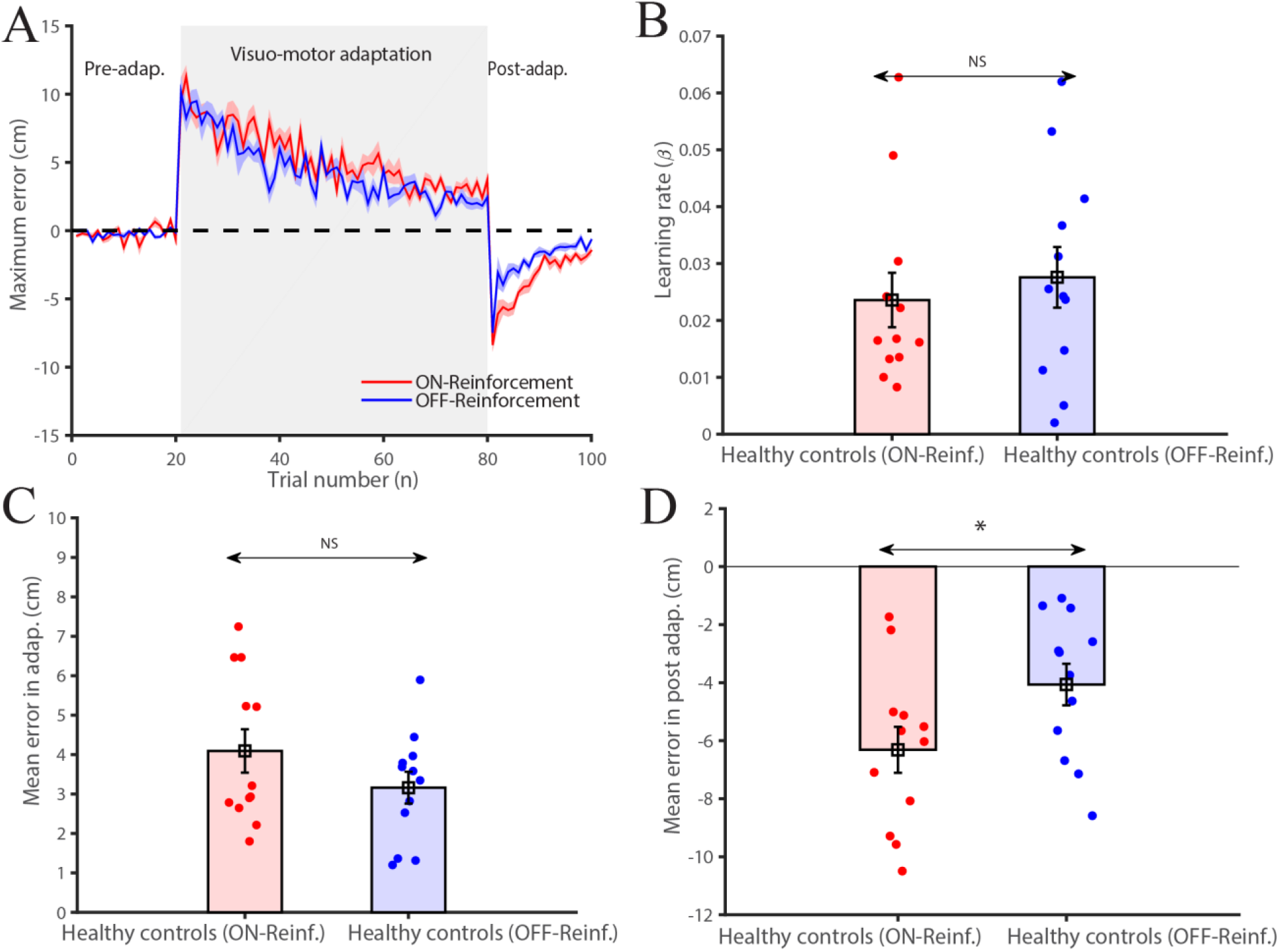
Healthy controls with and without reinforcement in visuomotor adaptation (A) Maximum error in pre-adaptation, visuomotor adaptation, and post-adaptation across healthy controls with reinforcement (n=20) and healthy controls without reinforcement. Red indicates healthy controls with reward and blue indicates healthy controls without reinforcement. (B) Learning rate differences in healthy controls with reinforcement (red) and healthy controls without reinforcement (blue) (n=20) reveal no differences in learning with and without reinforcement. (C) Averaged mean errors in healthy controls with reinforcement (red) was similar to healthy controls without reinforcement (blue) (n=20). (D) Averaged mean errors in healthy controls with reward (red) and healthy controls without reinforcement (blue) in postadaptation revealed greater error post-adaptation in healthy subjects with reinforcement.

In addition to the loss of dopaminergic input caused by the degeneration of neurons in substantia pars compacta, the progression of PD is also thought to reconfigure the connections within the basal ganglia (Albin *et al.*, 1989). To distinguish between these hypotheses, we also analyzed the correlation between the rate of change in the learning during OFF-and ON-states with the difference of the severity of motor symptoms measured by Unified Parkinson’s Disease Rating Scale part –III (UPDRS-III) during OFF-dopamine and ON states. We observed no correlation between the differences in the UPDRS-III scores in ON and OFF state with differences in the learning rate with or without reinforcement (Figure 7; r = 0.15, p = 0.63 (without reinforcement), r = 0.12, p = 0.60 (with reinforcement) and r = 0.26, p = 0.41 (DBS)). This analysis suggests that the difference in learning rates may reflect changes in dopamine, rather than motor adaptations in basal ganglia as a consequence of motor symptoms of PD.

**Figure 7:**
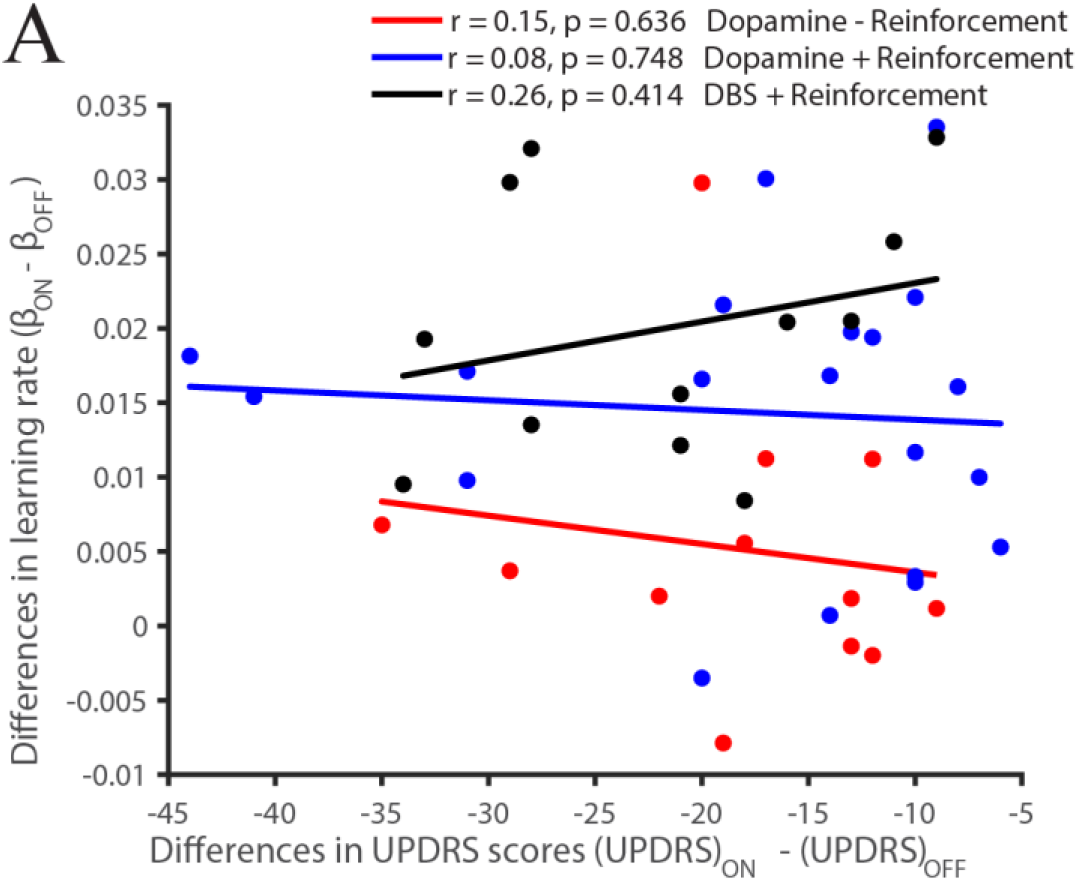
Correlation between differences in UPDRS scores with the differences in learning rate of Parkinson’s patients showed no correlation, without reinforcement (red), with reiforcement (blue) and with subthalamic deep brain stimulation (black).

## Discussion

In this study, we made two significant observations. First, we demonstrated how the presence and absence of dopamine and STN stimulation influenced the rate of adaptation, thereby implicating the role of basal ganglia in visuomotor adaptation. Second, we also showed that reinforcement at the end of the trial profoundly affected drug-induced (dopaminergic) learning in PD patients. Taken together, these results indicate a link connecting reinforcement, dopamine and basal ganglia in the modulation of visuomotor adaptation.

We examined visuomotor adaptation using a well-studied visuomotor perturbation (error-based task) with a few small modifications. Firstly, subjects had to learn to compensate for a rotation of 45° whereas in most previous work the rotations are typically 30° rotations. Secondly, the subjects made the movements on a table top but observed the effects on the screen. Thus one could argue that a larger error and the more complex motor to vision mapping may have involved basal ganglia by selectively engaging task-related errors (Desmurget *et al.*, 2004), preferentially engaging role of more explicit strategic components during learning (Taylor *et al.*, 2014; Mongeon, D. et al., 2013). Although, the current work was not designed to rule out the role of explicit learning, since patients with cerebellar degenerative diseases also exhibited similar deficits, we beleive that our task tapped into the standard visuomotor adaptation paradigm that also involves the cerebellum (Martin *et al.*, 1996; Maschke, 2003; Smith, 2005; Tseng *et al.*, 2007; Criscimagna-Hemminger *et al.*, 2010; Donchin *et al.*, 2012). In addition, the pattern of learning deficits was not restricted to the initial component of learning when the errors were large but rather reflects a global decrease in learning rate captured by the exponential fit, indicating that the basal ganglia contribution was not just restricted to the initial component of learning. This is in contrast to (Mongeon *et al.*, 2013) who showed that PD patients had a selective deficit for larger and possibly more explicit errors but not for smaller and presumably more implicit errors. Although the differences in our results compared to Mongeon et al 2013, as well as previous work Marinelli *et al.*, 2009; Bédard & Sanes, 2011; Semrau *et al.*, 2014 showing the absence of dopaminergic medication during visuomotor learning, are not clear; to the best of our knowledge this is the first study to use the same subjects in both the ON and OFF states. The ability to use the same subject as their own control has a major advantage given the complexities of the disease and subsequent alteration of the basal ganglia circuitry that may preclude using aged matched healthy subjects as a valid control; in addition to the obvious benefit of increased statistical power. However, a potential caveat is that transfer of learning within a patient could confound the interpretations of results. This was minimized by randomizing the order by which the groups were exposed to the adaptation task such that OFF and ON groups were counterbalanced. Furthermore, analysing the subjects separately based on the order in which ON and OFF patients were exposed to the task revealed the same result, precluding order as a potential confound in our analyses.

Another issue that previous studies did not address concerns whether dopaminergic projections from the ventral tegmental area could have also mediated these effects independent of the basal ganglia (Ikai *et al.*, 1992; Melchitzky & Lewis, 2000). Here we show for the first time that deep brain stimulation of the subthalamic nucleus (STN) produces similar effects on motor learning mimicking the dopamine ON and OFF condition. Although this study is agnostic about whether STN directly modulates cerebellar activity (Hoshi *et al.*, 2005; Bostan *et al.*, 2010), or whether its effects are mediated indirectly by modulating the level of striatal dopamine (Jenkinson & Brown, 2011; Min *et al.*, 2016), our findings firmly establish that the basal ganglia participate in visuomotor adaptation. In support of this view, a recent study by Brown and colleagues (Tan *et al.*, 2014) reported that low-frequency beta power from the STN is correlated with performance error in a visuomotor learning task indicating the role of basal ganglia in visuomotor adaptation. However, since this study did not assess motor learning in the presence and absence of stimulation, the causal contribution of STN could not be verified.

We propose that the ability to learn from errors is also dependent on the basal ganglia and is driven by reinforcement of successful actions. This hypothesis is congruent with prior results indicating that reinforcement can modulate visuomotor adaptation (Huang *et al.*, 2011; Galea *et al.*, 2015; Nikooyan & Ahmed, 2015; Kim *et al.*, 2018). Interestingly, in the context of PD patients such modulation is not observed during the learning of a new task but is typically observed as deficits in savings or relearning of a learnt rotation, which is compromised in PD patients (Marinelli *et al.*, 2009; Bédard & Sanes, 2011; Leow *et al.*, 2012). This is in contrast to our results which show modulations during learning. Thus, the mechanism by which reinforcement influences visuomotor adaptation is not clear. There are two potential mechanisms by which reinforcement learning can modulate visuomotor adaptation. In the first mechanism, a reinforcing signal from the basal ganglia can potentiate learning independently of cerebellum based learning (Huang *et al.*, 2011; Haith & Krakauer, 2013). Such a mechanism is fundamentally additive in nature. In the second arrangement, the reinforcing signal can interact with cerebellum based learning by modulating the sensitivity to errors (Galea *et al.*, 2015; van der Kooij & Overvliet, 2016). Such a mechanism is multiplicative in nature.

In this study, these hypotheses could be tested by manipulating the levels of dopamine (ON versus OFF) and secondary reinforcement delivered at the end of the trial (tone). Although such secondary reinforcement is unlike a primary reinforcement signal like food reward, we believe that subjects rapidly associate the delivery of the tone to successful performance through association. In this respect, primate studies have shown that dopamine neurons not only respond when the animal receives a reward but also when a stimulus such as a tone predicts a reward (Shultz et al 2016). Consistent with idea that the tone serves as a reinforcement signal, we observed a profound difference in learning in the presence and absence of the tone. Specifically, we observed a multiplicative interaction between these two variables such that the difference between OFF-dopamine and ON-dopamine in the presence of reinforcement, disappeared in the absence of reinforcement (see Figure 5). Thus, our results are most easily explained by a gain hypothesis in which reinforcement mediated at the end of the completion of a successful trial in the presence of releases dopamine in the proportion that changes the extent of learning occurring in the cerebellum. These differences in learning as a consequence of reinforcement do not appear to be an effect of decreased vigor (Hughes *et al.*, 1992) since we did not find any significant difference in the reaction times and disease state with and without reinforcement (see Table 5). Furthermore, we observed no difference in number of correct trials as a fraction of the average number of trials for each epoch for all groups and all experiments (see Table 6). However, increases in movement time are seen in controls and patients across all the conditions. This is not entirely surprising given that that the rotation imposes an error in the trajectory that needs to be corrected.

**Table 5:** Reaction time and movement time across groups

| Patients groups |  | Pre-adaptation | Visuo-motor adaptation | Post-adaptation |
| --- | --- | --- | --- | --- |
| Ataxia | RT | 685 ± 128 | 699 ± 99 | 626 ± 88 |
|  | MT | 1060 ± 150 | 1478 ± 112 | 1135 ± 113 |
| Ataxia group age matched healthy control | RT | 420 ± 40 | 441 ± 45 | 421 ± 39 |
|  | MT | 645 ± 54 | 991 ± 105 | 840 ± 73 |
| Parkinson's OFF-dopamine | RT | 573 ± 78 | 672 ± 85 | 625 ± 77 |
|  | MT | 976 ± 178 | 1465 ± 120 | 1061 ± 131 |
| Parkinson's ON-dopamine | RT | 578 ± 81 | 589 ± 84 | 550 ± 65 |
|  | MT | 818 ± 139 | 1286 ± 138 | 1006 ± 106 |
| Parkinson's group age matched healthy control | RT | 448 ± 40 | 465 ± 40 | 446 ± 40 |
|  | MT | 759 ± 99 | 1187 ± 126 | 895 ± 80 |
| Parkinson's OFF-deep brain stimulation | RT | 1028 ± 342 | 1115 ± 324 | 685 ± 214 |
|  | MT | 1315 ± 113 | 1588 ± 128 | 1264 ± 157 |
| Parkinson's ON-deep brain stimulation | RT | 835 ± 203 | 857 ± 200 | 786 ± 203 |
|  | MT | 1094 ± 219 | 1416 ± 95 | 1168 ± 96 |
| Parkinson's age matched healthy control | RT | 444 ± 56 | 468 ± 57 | 447 ± 58 |
|  | MT | 622 ± 87 | 1057 ± 153 | 876 ± 96 |
| Parkinson's OFF-dopamine without reinforcement | RT | 554 ± 111 | 636 ± 102 | 571 ± 71 |
|  | MT | 1075 ± 188 | 1365 ± 299 | 1273 ± 156 |
| Parkinson's ON-dopamine without reinforcement | RT | 508 ± 89 | 578 ± 124 | 528 ± 87 |
|  | MT | 930 ± 183 | 1462 ± 162 | 1175 ± 113 |
| Parkinson's age matched healthy control without reinforcement | RT | 464 ± 108 | 500 ± 112 | 505 ± 160 |
|  | MT | 693 ± 148 | 1006 ± 141 | 841 ± 127 |

**Table 6:**
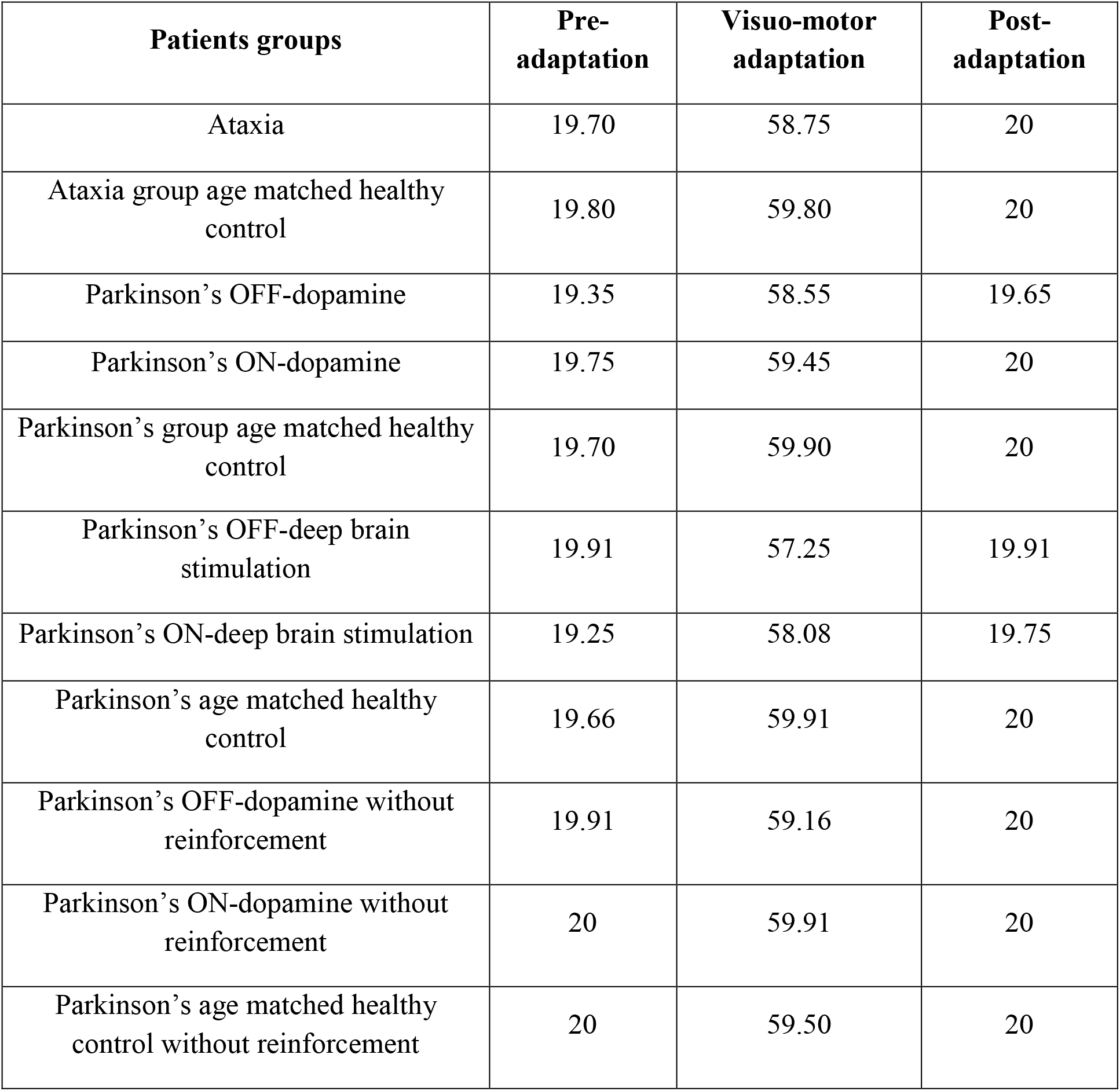
Mean number of correct trials across groups.

Surprisingly, while the presence and absence of reinforcement appeared to have a profound effect on learning in PD patients, learning was unaffected in healthy controls. This could indicate, other strategies such as aiming (Huang *et al.*, 2011; Taylor *et al.*, 2014), the use of redundancy (Singh *et al.*, 2016) and possibly the use of other factors such as increased co-contraction that may mitigate the effect of perturbation and may compensate for the lack of the effect of reinforcement in controls. Nonetheless, the post-adaptation after-effect did reveal the effect of reinforcement on controls as well. This differential effect could either indicate that post-adaptation after-effect may be a more sensitive measure of learning and or, may reflect a higher retention of motor memories due to reinforcement (Galea *et al.*, 2015). These results are consistent with recent work showing that rewards did not influence the rate of learning, but help in motor retention (Galea *et al.*, 2015), although the presence of visual feedback as well as the absence of an interval between adaptation and post-adaptation epochs in our task precludes a clear separation between after affects, unlearning and retention. In this context, it is also interesting to note that deep brain stimulation of the subthalamic nucleus (STN) produces similar effects on motor learning mimicking the dopamine ON and OFF condition but do not show clear post-adaptation after-effect. Whereas a clear after-effect was seen between dopamine ON and OFF conditions, suggesting different mechanisms at play for learning during the adaptation and post-adaptation epochs. Taken together, we interpret these results as indicating, a specific role for dopamine/STN from the basal ganglia in modulating learning during visuomotor adaptation, while post adaptation after-effects may be mediated by dopamine but through a mechanism that may not involve the basal ganglia but may involve direct dopaminergic projections to the cerebellum from the ventral tegmental area (Ikai *et al.*, 1992; Melchitzky & Lewis, 2000).

## Acknowledgment

We thank Prof. Dibakar Sen for lending us the portable Polhemus System and Dr. Sumitash Jana for the initial setup of the experiment. Dr. Ravi Yadav and Dr. Dwarakanath Srinivas for help in DBS patients recording work.

